# Vitreous Inflammation as an Early Indicator of Retinal Ganglion Cell Loss Following Acute Optic Nerve Injury in Mice

**DOI:** 10.64898/2025.12.20.694725

**Authors:** Shichu Chang, Weijia Fan, Jiahui Wu, Lei Xu, Virginia Lee, Charlotte Zhang, Edward P. Kronenberg, Wenjin Xu, Hu Yang, Hao F. Zhang, Xiaorong Liu

## Abstract

Retinal ganglion cell (RGC) degeneration in optic neuropathies is often preceded by neuroinflammatory changes, yet the earliest *in vivo* indicators of this process remain poorly defined. Here, we identified vitreous hyperreflective foci (VHRFs) by visible-light optical coherence tomography (vis-OCT) as an early inflammatory signature of RGC injury. Longitudinal vis-OCT imaging after optic nerve crush (ONC) revealed that VHRFs emerged as early as 6 hours post-injury and peaked before the significant RGC loss. Confocal analysis showed that these VHRFs corresponded to activated amoeboid microglia undergoing vertical migration from the outer to inner retina and horizontal movement toward the optic nerve head area. Similar amoeboid microglia were also observed in the anterior segment, suggesting a global ocular inflammatory response to the ONC injury. RNAscope *in situ* hybridization further demonstrated elevated *IL-1β* expression in vitreous amoeboid microglia. Moreover, pharmacological blockade of IL-1 signaling with an IL-1 receptor antagonist significantly reduced VHRFs, suppressed microglial migration, and delayed RGC loss. Taken together, our findings identify VHRFs as a previously unrecognized early danger signal for RGC degeneration and highlight IL-1–mediated inflammation as a tractable early therapeutic target for preventing RGC degeneration and vision loss.

**Graphical Overview:** 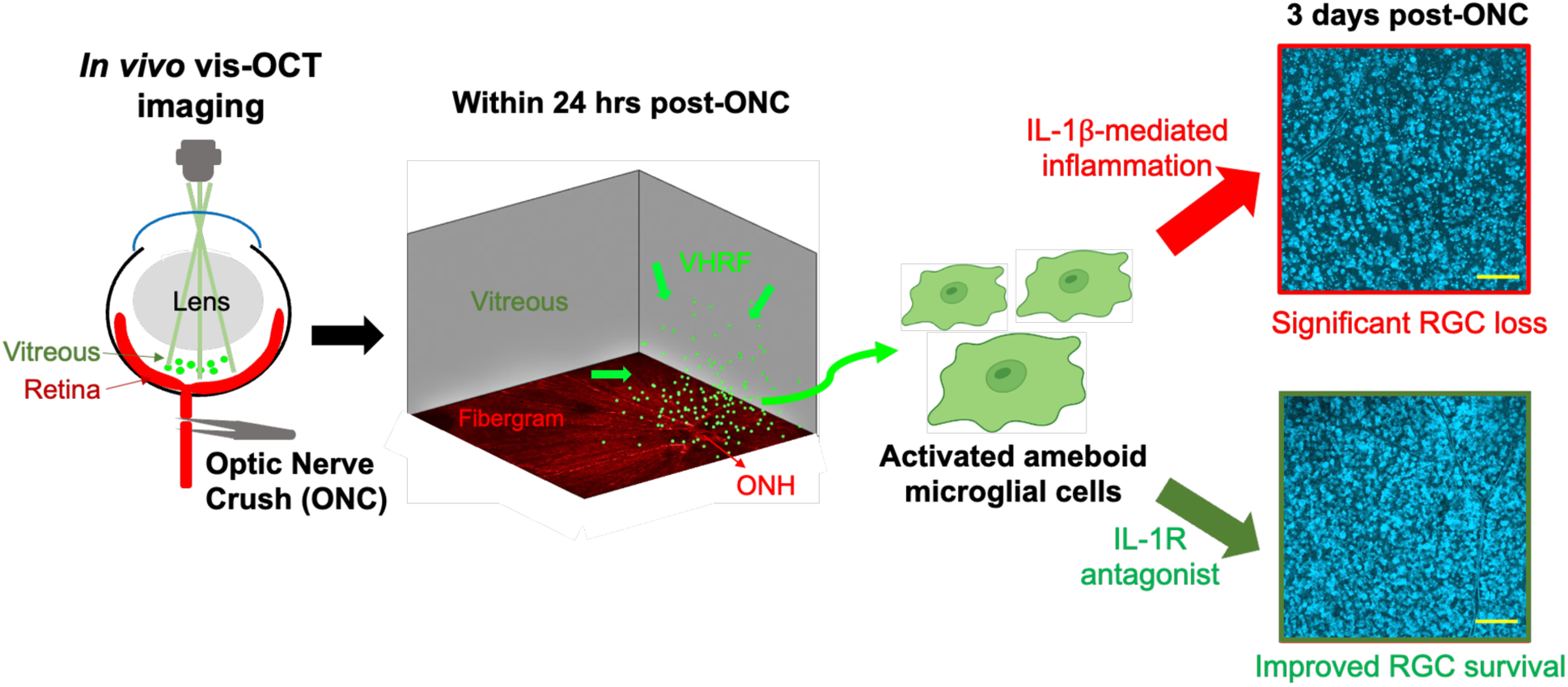

## Introduction

Retinal ganglion cells (RGCs) are the output neurons of the retina, and their axons form the optic nerve that conveys visual information to the brain [1]. RGC degeneration is a hallmark of many different ocular and neurodegenerative diseases, leading to irreversible vision loss [2]. The underlying insults can be acute, such as traumatic [3] and ischemic optic neuropathy [4], and optic neuritis [5], or chronic, as seen in glaucoma [6] and neurodegenerative disorders (*e.g.*, Alzheimer’s Disease, Parkinson’s Disease, Huntington’s Disease) [7,8]. Because RGC death is irreversible, early detection and intervention are crucial for preserving RGCs and vision [9–11]. Despite significant advances in structural and functional imaging, existing biomarkers often fail to capture the earliest events leading to RGC degeneration [12–14]. In other words, there remains an urgent need for highly sensitive imaging approaches and specific early biomarkers capable of detecting RGC distress before irreversible damage occurs [15].

Optical coherence tomography (OCT) is a noninvasive *in vivo* imaging technique that has been widely applied in both research and clinical settings [16]. We established a novel visible-light OCT (vis-OCT) imaging system with an improved axial resolution, enabling the detection of subtle differences in the retinal layer structures in mouse [16–19], tree shrew [20,21], and human eyes [21–23]. Using this system, we recently detected a subtle vitreous signal, termed hyperreflective foci (HRFs), that emerged after ONC and had not been characterized by conventional OCT methods [24]. Indeed, HRFs were observed in many inflammation-related ocular diseases, such as diabetic retinopathy (DR) [25], age-related macular degeneration (AMD) [26], uveitis [27], and macular edema [28]. They were identified as melano-macrophages in post-mortem samples from AMD patients [26], and are commonly believed to be inflammatory cells in uveitis [29]. In ONC and glaucoma models, some studies suggest the early activation of microglial cells and astrocytes before substantial RGC loss in the mouse retina or optic nerve [30–33]. However, the molecular and cellular identity of the vitreous HRFs (VHRFs) and the roles they play in RGC loss remain to be investigated.

With early detection, timely intervention is equally critical for protecting RGCs. However, the precise molecular triggers to RGC loss are not well defined despite extensive investigations on disease mechanisms [34–36]. Studies have shown that multiple pathological processes are involved in early RGC loss, including oxidative stress [37], mitochondrial dysfunction [38], neurotrophic factor deprivation [39], and neuroinflammation [40]. Among these, neuroinflammation emerged as a promising early danger signal and therapeutic target. For example, at only 12 hours post-ONC, the vitreous myeloid cells were significantly elevated [24]. Within 24 hours post-ONC, retinal microglia activation, evidenced by increased expression of NLRP3 inflammasome and tumor necrosis factor-α (TNFα), was also detected [31,41]. At 24 hours post-ONC, the glial fibrillary acidic protein (GFAP) marker for astrocytes and Müller cells also significantly increased in injured retinas, indicating macroglia activation [42]. Similarly, in glaucoma models, microglia and macroglia activation were also detected preceding massive RGC loss [32,43–45]. In response to major retinal insults, pro-inflammatory cytokines affect neuronal survival. In the retina, studies have indicated that inhibition of pro-inflammatory pathways, including IL-1 receptor antagonism [46], IL-6 knockout [47], and IL-1α/ TNFα/ C1q triple knockout [48], can be protective to RGCs. Therefore, in this study, we first tracked the inflammatory signal of VHRFs *in vivo* by vis-OCT imaging post-acute ONC injury and then determined their cellular identity. Next, we perturbed the activation and mobilization of VHRFs through the IL-1 signaling to assess their impact on RGCs.

## Methods

Healthy adult (3–6 months) male and female wildtype C57BL/6 mice were used for this study. The CX3CR-1^GFP^ knock-in mice with C57BL/6J background (The Jackson Laboratory, 005582) were also used. All animal protocols were approved by the University of Virginia institutional animal care and use committee (IACUC) and complied with the guidelines of National Institutes of Health (NIH) and the Association for Research in Vision and Ophthalmology (ARVO).

### Optic Nerve Crush Surgery and Drug Administration

The ONC surgery was performed as previously described [18,19,31]. In brief, mice were first anesthetized with intraperitoneal injection of a mixture of ketamine (100 mg/kg, Kataset, Zoetis, Parsippany-Troy Hills, NJ, USA; NADA no. 043-304) and xylazine (8 mg/kg, AnaSed, Akorn Pharmaceuticals, Gurnee, IL, USA; NADA no. 139-236). In the temporal and superior part of the conjunctiva, a small incision was made, and the optic nerve was exposed by blunt dissection of the surrounding muscles. The optic nerve was then clamped with Dumont #5/45 Forceps (Fine Science Tools, Foster City, CA, USA; No. 11251-35) at around 1 mm behind the eyeball for 8-10 seconds.

Right after the ONC surgery, a pair of forceps was used to hold the conjunctiva to stabilize the eyeball, and a 2.5 μL Hamilton syringe (Hamilton REF#7632-01, 62RN, USA) mounted with 26-gauge needle (Hamilton REF#7758-02) was applied to penetrate the cornea. 40 or 20 μg of IL-1Ra Anakinra (20 mg/mL in PBS, MedChemExpress, HY-108841) or PBS with the same volume was injected into the anterior chamber of the eye. Both eyes of the same animals were crushed, with one eye injected with PBS and another injected with Anakinra. After injection, moxifloxacin (0.5%, NDC 60505-05824; Apotex Corp., Toronto, Canada) was applied to the injected eyes to prevent infection. Mice were kept on a heating pad until they fully recovered.

### Vis-OCT Imaging and Data Processing

Mice were first anesthetized with intraperitoneal injection of a mixture of ketamine (114 mg/kg, Kataset; Zoetis) and xylazine (17 mg/kg, AnaSed) to induce stabilized anesthesia for 30-40 min. The pupils were dilated with tropicamide drops (1%; Henry Schein Animal Health, Covetrus, Portland, ME, USA). During imaging, mice were kept warm with an infrared heat lamp and given artificial tear drops (1.4%; Rugby Laboratories, Inc., Hempstead, NY, USA) to prevent corneal dehydration.

Mice were imaged using a small-animal vis-OCT system (Halo 100; Opticent Health, Evanston, IL, USA) as previously reported [17,18]. For each eye, a single volumetric scan was acquired with the optic nerve head (ONH) positioned in the corner of the field of view (FOV) to capture both the ONH and surrounding retinal regions. Each volume consisted of 512 B-scans and 512 A-lines per B-scan, covering a 547 μm × 547 μm area. The system operated at 40 kHz, with each volume acquired in approximately 10.5 seconds.

Using a previously published MATLAB code [24], vitreous regions were extracted from each acquired volume for visualization of vitreal hyperreflective foci (VHRFs). The inner retinal surface in each B-scan was segmented based on its intensity gradient, and 11 to 227 μm above this boundary was defined as the vitreous. Contrast enhancement and normalization were subsequently applied to the extracted vitreous regions. VHRFs were detected using an intensity threshold (ten times the standard deviation from the mean intensity) followed by a size threshold (105.1 μm^3^ < signal volume < 21021.3 μm^3^) as small, bright, spherical spots on the vis-OCT images. Finally, the total number of VHRFs within this 547 μm × 547 μm × 216 μm volume was quantified, and the volumetric VHRF density (#VHRFs/mm^3^) from each vitreous volume was calculated.

### Immunohistochemistry and Confocal Imaging

Mice were euthanized with euthasol (600 mg/kg, Euthasol, Virbac ANADA, no. 200-071; Virbac, Carros, France) and perfused with phosphate-buffered saline (PBS) and then 4% paraformaldehyde (PFA, ChemCruz, sc281692; Santa Cruz Biotechnology, Dallas, TX, USA). After perfusion, the eyes were enucleated.

For whole-eye sectioning, enucleated eyes were soaked in 4% PFA followed by 20% sucrose overnight. They were then embedded into optimal cutting temperature (OCT) compound and cryo-sectioned at 35-μm thickness sagittally to preserve the vitreous humor. For retinal flat mounts, the eye cups were dissected, post-fixed in paraformaldehyde for 30 minutes, washed with PBS 3 times, and PBS containing Triton-X detergent (PBST, 0.5% Triton X-100) 1 time. Both whole-eye sections and eye cups were blocked for one hour at room temperature in blocking buffer (1% BSA and 10% normal donkey serum, 0.5% Triton X-100, Sigma-Aldrich, St. Louis, MO, USA).

Primary antibodies, including anti-Iba1, GFAP, CD68, CD34, Ki67, and Laminin (Table 1), were applied to the sections, overnight at 4°C; anti-Rbpms (Table 1) was applied to eye cups, overnight at 4°C [19]. Secondary antibodies, including donkey anti-rabbit immunoglobulin G conjugated to Alexa Fluor 647 dye (A-31571, RRID: AB_162542; Invitrogen) and donkey anti-mouse immunoglobulin G conjugated to Alexa Fluor 594 dye (A-21203, RRID: AB_141633; Invitrogen) were diluted at 1:1000 in blocking buffer and incubated overnight at 4°C. After immunostaining, the whole-eye sections were directly cover-slipped, while retinas in eye cups were cut into four quadrants and flat-mounted on the slides before being cover-slipped.

**Table 1.**
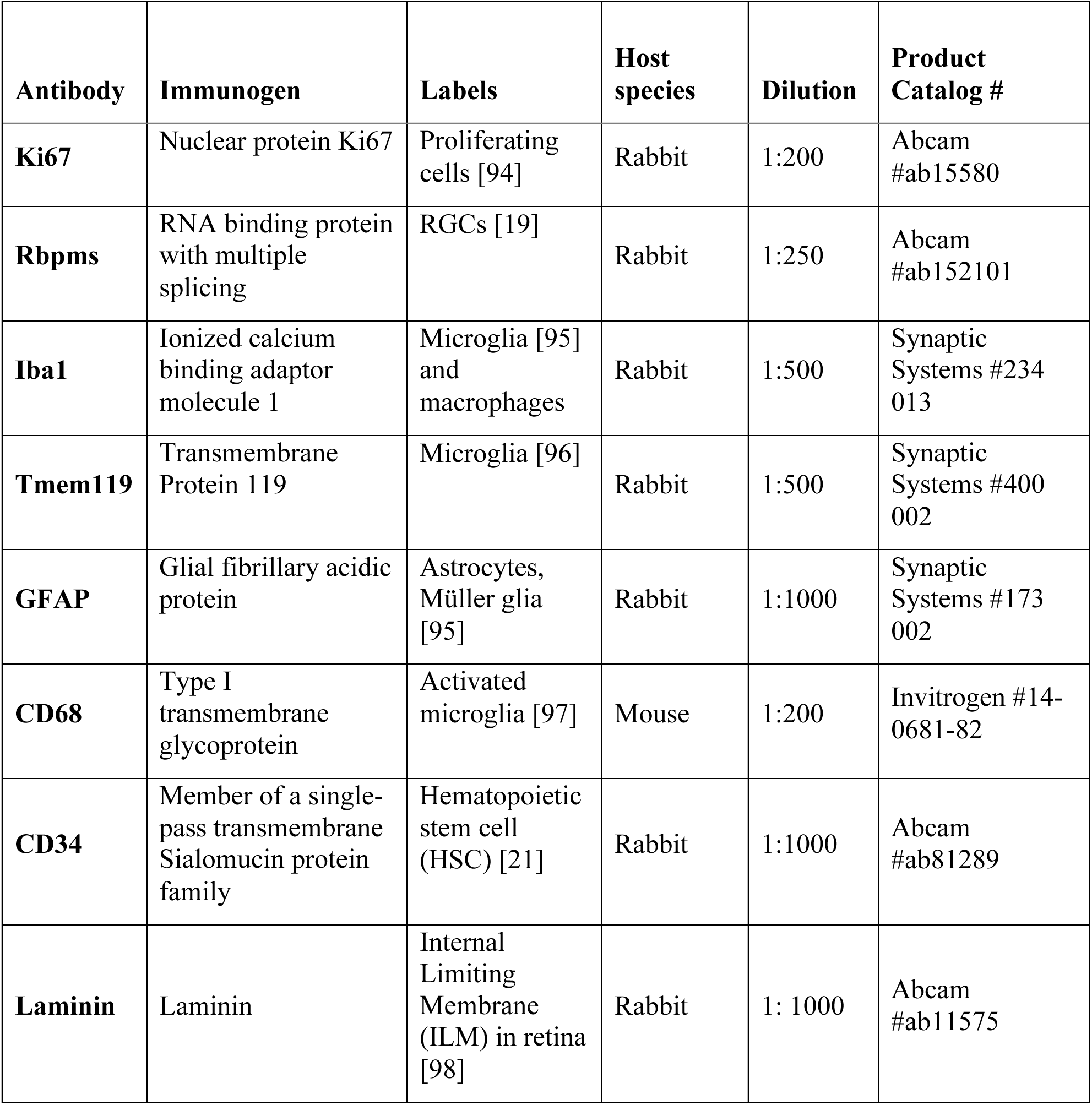
Primary Antibody list.

Confocal images were taken using a Zeiss LSM800 confocal microscope (Zeiss, Thornwood, NY, USA). For retinal flat mounts, z-stack images covering the depth of the outer nuclear layer to the GCL (approximately 50–80 μm) were acquired. Individual images were captured using the tiling function in Zen (Zen 3.8; Zeiss, Oberkochen, Germany) at 10× magnification, covering the entire retina. For whole-eye sections, z-stack images covered the depth of the entire 35-μm thickness, and the tiling function was applied to cover the area of the lens, vitreous, and retina.

### RNAscope *in situ* hybridization

RNAscope *in situ* hybridization was performed using the RNAscope Multiplex Fluorescent Detection Kit V2 (ACD Bio, Newark, CA, USA). All tools, containers, and desktops were cleaned with RNaseZAP (Sigma-Aldrich, St. Louis, MO, USA; R2020) to prevent RNA degradation. 20 μm retina sections were post-fixed in 4% PFA for 2 hours and washed in PBS for 5 min. Next, the sections were treated with RNAscope Hydrogen Peroxide for 10 min, and then submerged in dH_2_O for 5 min. After dehydration with 50%, 70% and 100% ethanol for 5 min each, the sections were dried in a 60°C oven for 30 min, then kept at room temperature overnight.

On the second day, after drawing a hydrophobic barrier around sections, protease III was applied for 30 min at 40°C. Next, the sections were incubated with target probes (Table 2) for 2 hours at 40°C and then treated with amplification reagents at 40°C (AMP1 for 30 min, AMP2 for 30 min, and AMP3 for 15 min). Afterwards, TSA Fluorescein, Cy3, and Cy5 fluorophores were applied to sections at 40°C [49]. Between each 40°C treatment step, the sections were submerged in wash buffer for 2 × 2 min at room temperature. Finally, mounting medium with DAPI (VECTASHIELD, Newark, CA, USA, H-1200-10) was applied, and the slides were cover-slipped for confocal imaging.

**Table 2.** RNAscope Probe list.

| **Probe Name** | **Source** | **Identifier** | **Labels** |
| --- | --- | --- | --- |
| RNASCOPE® PROBE -<br>MM-IL1B-C3 | ACD | Cat# 316891-C3 | IL-1beta |
| RNASCOPE® PROBE -<br>MM-TNFA | ACD | Cat# 311081 | TNF-alpha |

### Cell Quantification and Statistical Analyses

For RGC soma quantification, we followed previously published protocols [19,50,51]. Briefly, mouse retinas were immunostained with anti-rbpms antibody. For each retina, 18-22 enface z-stack images covering the depth of the GCL (20–30 μm) were captured. On each quadrant of the retina, 3 rectangles covering 0.03 mm^2^ area were randomly drawn on the images without overlapping with large blood vessels. All rbpms+ cells within the rectangles were manually counted in Zen. Cell density was calculated by total cell counts divided by area, and the cell densities of all 12 regions in one retina were averaged. For microglia quantification, CX3CR-1+ cells in a whole-eye section/ specific layers of the retina (e.g., RNFL+GCL, IPL, and OPL) were manually counted in Zen. All data analyses were conducted by two independent experimenters, and the average percentage difference between the two sets of data is 2.29 %.

All statistical analyses were performed using MATLAB (MathWorks, Natick, MA, USA) and Prism 10 (Dotmatics, Boston, MA, USA). For comparison between two groups of conditions (e.g., control and ONC comparison), we performed Student’s *t*-test. For comparison between three or more conditions (*e.g.*, multiple time points post-ONC, multiple antibody staining, and multiple drug treatment), we performed one-way analysis of variance (ANOVA) followed by Dunnett’s test for multiple comparisons. For the comparison between two independent sets of conditions (e.g., multiple time points post-ONC plus multiple retinal layers/ regions), we performed two-way ANOVA followed by Bonferroni’s multiple comparisons test. All results were reported as mean ± standard deviation.

## Results

### *In Vivo* Tracking of the Same Mouse Eyes Showed the Vitreous Signals Peaked within 24 Hours Post-Optic Nerve Crush

Using the vis-OCT imaging system that we established, we extracted the vitreous and the retinal nerve fiber layer (RNFL) signals as shown in Fig. 1A-B. We defined the vitreous area as a 547 μm × 547 μm × 216 μm volume above the retina and then quantified the total number of vitreous hyperreflective foci (VHRF) per eye (Fig. 1C). In our previous study, we analyzed vis-OCT data at a single time point across different mice [24]. To better characterize VHRF dynamics *in vivo*, here we longitudinally tracked signal changes in the same eyes following ONC injury. Our results showed that the density of the VHRFs began to emerge about 6 h post-ONC, peaked at 24 h, and started to decrease after 24 h (Fig. 1D). At 12 hours, VHRF density significantly increased from 100.8 ± 49.5 to 758.3 ± 394.0 VHRFs/mm^3^ (n = 6 eyes, p < 1e-4, One-way ANOVA/Dunnett’s post-hoc test, same below). The signal remained high at 24 hours post-ONC (853.5 ± 222.0 VHRFs/mm^3^, n = 6 eyes, p < 1e-4). The VHRF density gradually decreased to 436.5 ± 148.1 VHRFs/mm^3^ (n = 5 eyes, p = 5.5e-2) at 3 days post-ONC and 413.4 ± 111.9 VHRFs/mm^3^ (n = 4 eyes, p = 0.11) at 6 days post-ONC (Fig. 1E). In the sham operation group, there were very low background VHRFs compared with the baseline. The VHRF density changed from 75.6 ± 18.9 VHRFs/mm^3^ before sham operation to 75.6 ± 25.0 VHRFs/mm^3^ (n = 3 eyes, p > 0.99) at 12 hours, and 62.99 ± 21.82 VHRFs/mm^3^ (p = 0.93, n = 3 eyes) at 24 hours, suggesting the signals were not induced by surgical damage (Fig. 1E).

**Fig. 1.**
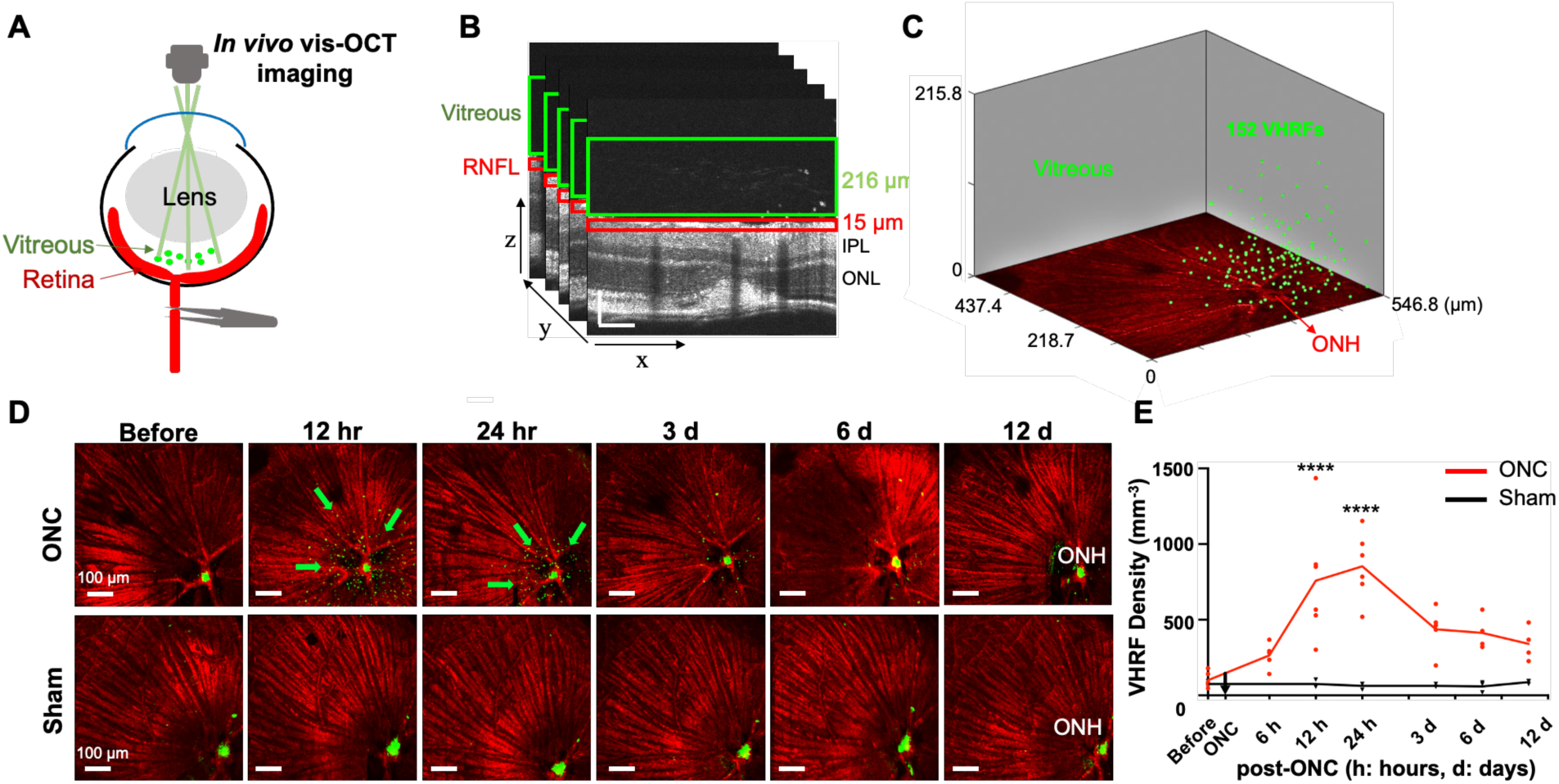
*In vivo* tracking of the same mouse eyes showed the vitreous signal peaked within 1 day post-ONC. (A) *In vivo* vis-OCT imaging. Visible light illumination (480 nm ∼ 650 nm) was delivered to the anesthetized mouse eye and reached the vitreous and retina. (B) Image processing for the vitreous and retina signals. From 11 to 227 μm above the internal limiting membrane (ILM) was segmented as the vitreous, and the superficial 15 μm of the retina was segmented as the retinal nerve fiber layer (RNFL). IPL: inner plexiform layer. ONL: outer nuclear layer. Scale bar: 100 μm. (C) 3-D localization of the vitreous signals at 12 hours post-ONC. (D) *In vivo* longitudinal tracking of the vitreous inflammatory signals. The fibergram was pseudo-colored red, and the vitreous signals were pseudo-colored green. Green arrows indicated VHRFs. (E) Changes of the inflammatory signal number with time in ONC and sham groups. ****: p < 0.0001 (One-way ANOVA/Dunnett’s post-hoc test).

Moreover, we monitored the VHRF dispersion over time (S-Fig.1A). Within 200 μm from the ONH, 3 days post-ONC (59.5 ± 20.5 %/mm^3^) showed significantly lower percentages of VHRF than 12 hours (90.5 ± 7.1 %/mm^3^, n = 5 eyes, p = 1e-4, Two-way ANOVA/Bonferroni’s post-hoc test, same below). Beyond 400 μm from the ONH, 3 days post-ONC vitreous (19.8 ± 15.3 %/mm^3^) had significantly higher VHRF percentage than 12 hours (1.6 ± 2.3 %/mm^3^, n = 5 eyes, p = 3e-2), suggesting the dispersion of VHRFs from the central ONH to more peripheral areas with time (S-Fig.1B).

### Vitreous Signals Preceded a Significant RGC Loss, Potentially Serving as an Early Danger Sign

We next determined the time course of vitreous signals and RGC loss post-ONC by immunostaining and confocal imaging (Fig. 2A, B). At 12 hours post-ONC, vitreous signal density had increased significantly (820.6 ± 301.3 VHRFs/mm^3^, n = 7 eyes) compared with baseline (100.8 ± 49.5 VHRFs/mm^3^, n = 6 eyes, p < 1e-4), whereas the RGC loss is not yet detectable (Ctrl: 4287 ± 188 cell/mm^2^, n = 5 retinas; ONC-12h: 4485 ± 325 cell/mm^2^, n = 4 retinas; p = 0.47). At 24 hours post-ONC, a significant RGC loss was first detected (3877 ± 237 cell/mm^2^, n = 5 retinas, p = 3.9e-2), immediately after the VHRF peak. The vitreous signals gradually died down at 12 days (340.1 ± 111.0 VHRFs/mm^3^, n = 4 eyes, p = 0.27), but, at the same time, the RGC loss became more significant (1015 ± 84 cell/mm^2^, n = 3 retinas, p < 1e-4, Fig. 2C, D). Together, our results show that vitreous signals emerge before significant RGC loss, suggesting their potential as early “danger” indicators of the upcoming neuronal loss.

**Fig. 2.**
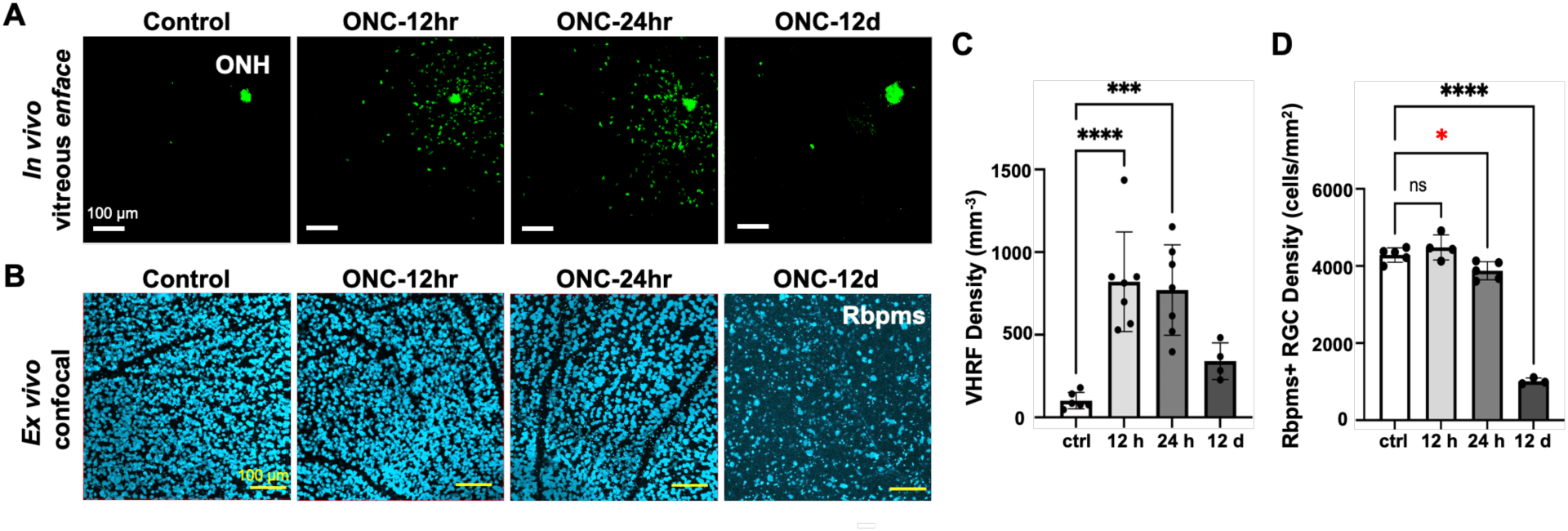
The vitreous signals preceded a significant RGC loss, potentially serving as an early “danger” sign. (A) Vitreous *enface* images at different time points of 4 different mice post-ONC. (B) Confocal images of flat-mounted retinas immunostained by Rbpms for quantification of RGC loss. (C) VHRF density at different time points post-ONC. Significant VHRF density increase was first detected at 12 and 24 hours. ***: p < 0.001; ****: p < 0.0001 (One-way ANOVA/Dunnett’s post-hoc tests). (D) Rbpms+ RGC density at different time points post-ONC. Significant RGC loss was first detected at 24 hours, but not at 12 hours. *: p < 0.05; ****: p < 0.0001 (One-way ANOVA/Dunnett’s post-hoc tests).

### The Vitreous Cells were Activated Amoeboid Microglia

We performed *ex vivo* whole-eye sectioning to characterize the VHRFs (Fig. 3A). We used the CX3CR-1^GFP^ knock-in line in which green fluorescent protein (GFP) is expressed in retinal microglia and macrophages. We found that the vitreous of ONC eyes exhibited markedly higher GFP than that of control eyes. Some vitreal signals were CX3CR-1 positive cells, co-labeled with ionized calcium-binding adaptor molecule 1 (Iba1, a marker for microglia), and glial fibrillary acidic protein (GFAP, a marker for astrocytes and Müller glia, Fig. 3B-C). We quantified the percentage of vitreous cells labeled by those markers and showed that 69.3 ± 11.8 % are CX3CR-1 positive among all the vitreous cells (DAPI+), 60.4 ± 10.0 % are Iba1 positive, and only 8.4 ± 2.4 % are GFAP positive (Fig. 3D). These results suggest that the vitreous signals are microglia but not Müller glia or astrocytes.

**Fig. 3.**
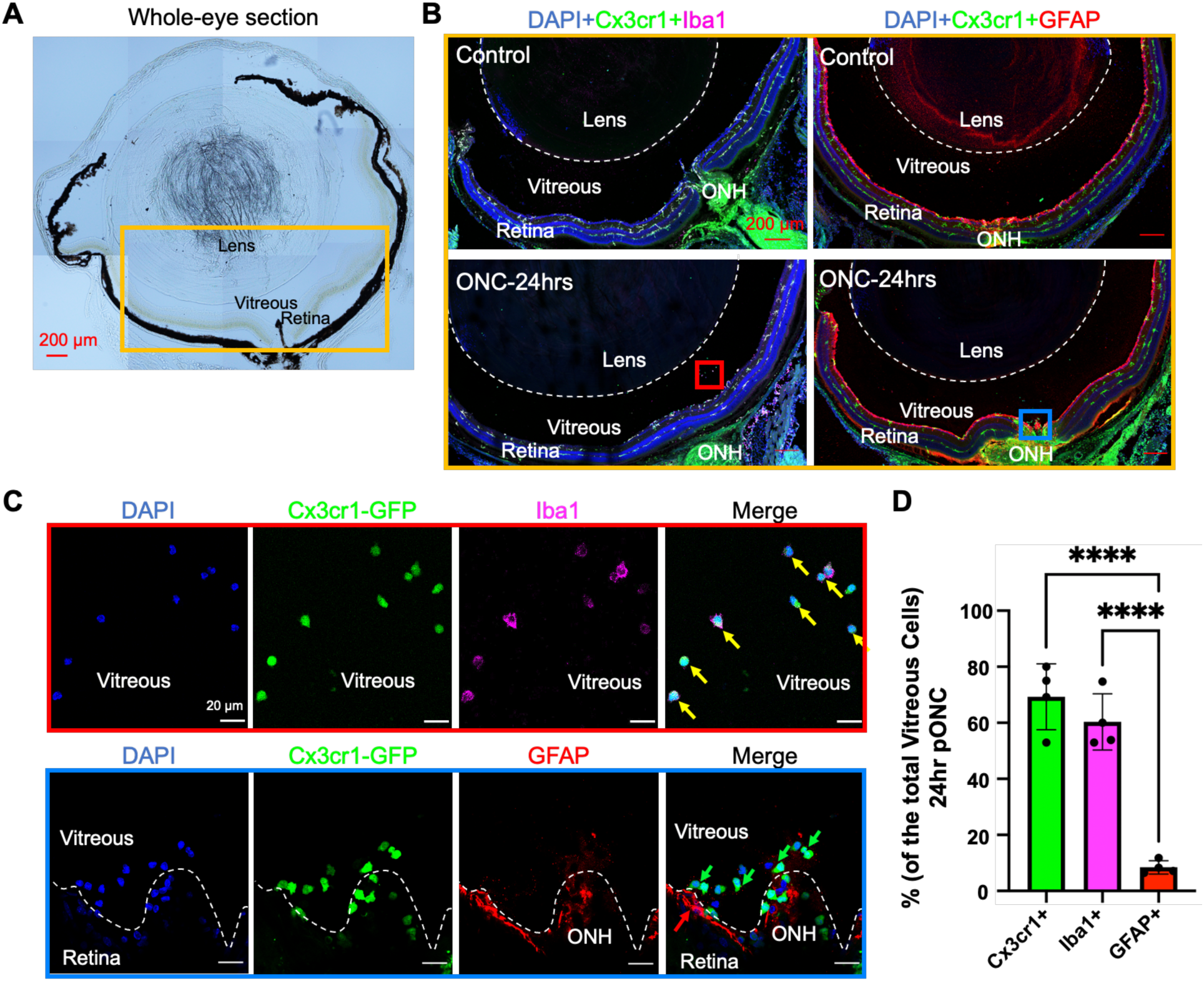
The Vitreous Cells were Microglia but not Müller Glia or Astrocytes. (A) Brightfield image of a whole eye section. Vitreous was preserved between the lens and retina. The orange box indicates the region imaged in panel B. (B) Iba1 and GFAP staining of control and 24 hours post-ONC CX3CR-1^GFP^ knock-in mouse eyes. ONH: Optic Nerve Head. (C) High magnification images of squared regions in panel B. Yellow arrows indicate CX3CR-1+Iba1+ cells; green arrows indicate CX3CR-1+ cells; red arrows indicate GFAP+ cells. (D) The percentage of CX3CR-1+, Iba1+, and GFAP+ cells among all vitreous cells (DAPI+). n = 4 mice. ****: p < 0.0001 (One-way ANOVA/Dunnett’s post-hoc test).

In addition, we noticed the distinct morphology of these VHRFs. The vitreous microglia adopted an amoeboid shape with no processes, different from ramified microglia at the resting state and activated microglia with enlarged cell bodies and shortened processes (Fig. 4A). In control retinas, the amoeboid microglia barely existed (2.3 ± 2.1 cells, n = 4 eyes), but at 24 hours post-ONC, their number significantly increased (39.2 ± 19.1 cells, n = 5 eyes, p = 6.7e-3, Student’s *t*-test, same below, Fig. 4B). These amoeboid microglia were observed to cross the internal limiting membrane (ILM, stained by anti-laminin) without causing detectable damage to the ILM (Fig. 4C).

**Fig. 4.**
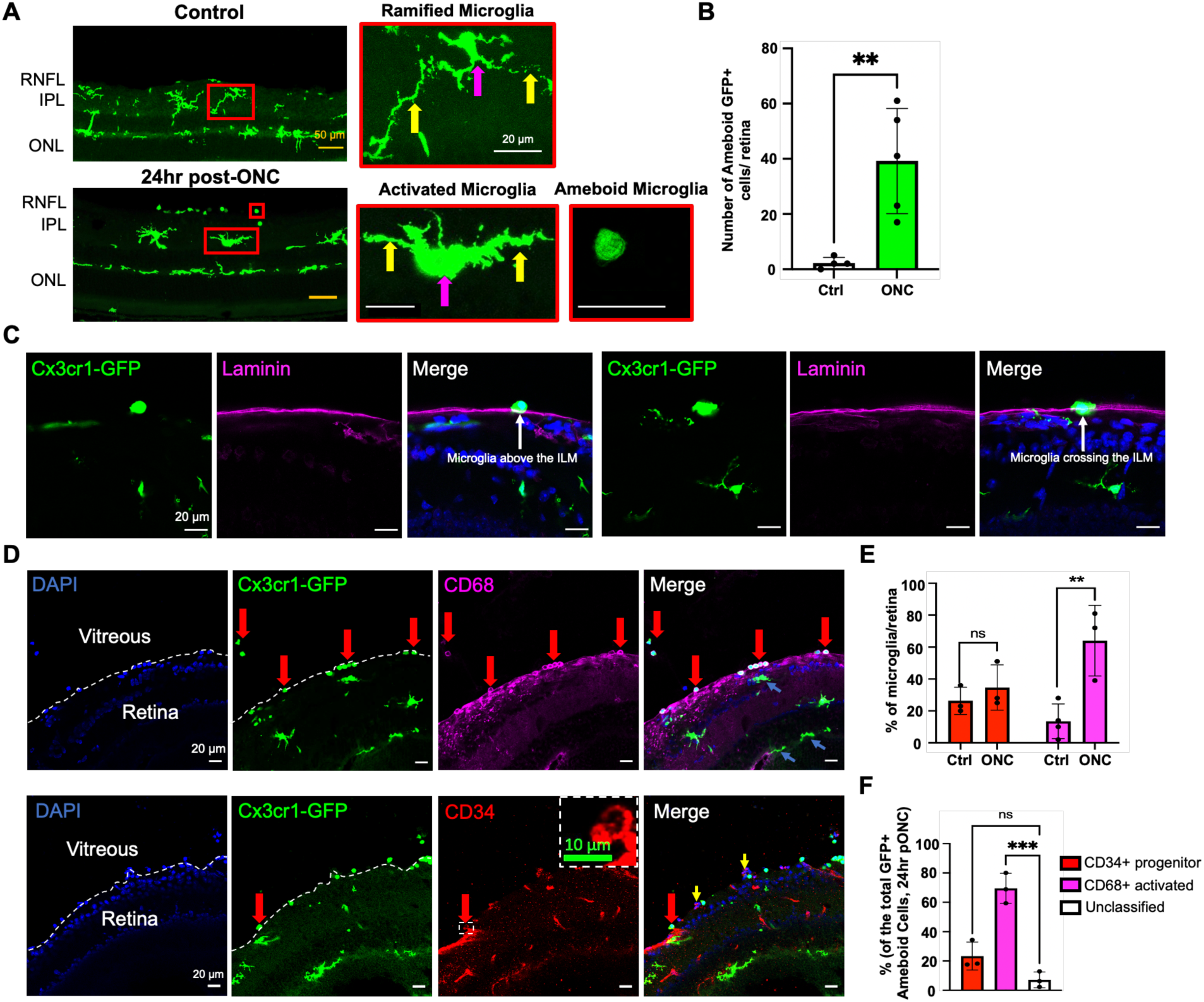
The majority of amoeboid microglia were activated microglia, while a small part of them were microglia progenitors. (A) Morphology of ramified, activated, and amoeboid microglia. Violet arrows indicate the cell bodies; yellow arrows indicate the processes. (B) Number of amoeboid microglial cells in control and ONC retinas. n = 4 mice. **: p < 0.01 (Student’s *t*-test). (C) Amoeboid microglia cross the internal limiting membrane (ILM) in the retina. ILM was immunolabeled by laminin antibody. White arrows indicate retinal amoeboid microglia. (D) CD34 and CD68 staining of CX3CR-1^GFP^ knock-in mice at 24-hr post-ONC. Red arrows indicate co-labeled cells; yellow arrows indicate CD34+ CX3CR-1-cells; blue arrows indicate CX3CR-1+ CD68-cells. (E) Number of CD34+ and CD68+ cells in ctrl and ONC retinas. **: p < 0.01 (Student’s *t*-test). (F) Percentage of CD34+ and CD68+ cells among CX3CR-1+ amoeboid microglial cells. ***: p < 0.001 (One-way ANOVA/Dunnett’s post-hoc test).

To examine whether the vitreous cells are likely to be activated microglia or microglia progenitors, we applied CD68 (a marker for activated microglia) and CD34 (a marker for microglia progenitors) on the whole eye sections (Fig. 4D). Compared to the controls (13.5 ± 10.9 %, n = 4 eyes), ONC retinas showed a significant increase in the CD68+ microglia (64.0 ± 22.1 %, n = 3 eyes, p = 9.8e-3) but not the CD34+ microglia (control: 26.3 ± 8.5 %, n = 3 eyes; ONC-24h: 34.7 ± 14.2 %, n = 3 eyes, p = 0.43, Fig. 4E). Among the total population of CX3CR-1 GFP+ amoeboid microglia, 69.4 ± 10.3 % are CD68+ activated microglia, and 23.4 ± 9.5 % are CD34+ progenitors (Fig. 4F). Indeed, we found some progenitor cells undergoing mitosis as indicated by their dividing nucleus and Ki67 staining (S-Fig.2C, D). Together, our data suggest that most of the vitreous signals are activated amoeboid microglia, and some are microglial progenitors.

### Retinal Microglial Cells were Mobilized from the Retina and Dispersed throughout the Eye

We next investigated whether the vitreous microglial cells originated from the retina. We examined the distribution and dispersion of microglial cells across different retinal layers and regions. First, the percentage of microglia in the RNFL+GCL significantly increased at 24 hours post-ONC (40.0 ± 2.7 % vs 22.0 ± 6.8 %, n = 4 retinas, p < 1e-4, Two-way ANOVA/Bonferroni’s post-hoc test, same below), while the percentage of microglia in the outer plexiform layer (OPL) significantly decreased at both 12 and 24 hours post-ONC (control: 42.8 ± 3.9 %; ONC-12h: 33.1 ± 4.3 %, p = 4.2e-2; ONC-24 h: 32.5 ± 4.4 %, p = 2.6e-2, n = 4 retinas, Fig. 5A, B). The percentage of microglia in the inner plexiform layer (IPL) also showed a slight increase at 12 h post-ONC (43.6 ± 4.0 % vs 35.2 ± 3.2 %, n = 4 retinas, p = 0.11), suggesting that overall microglial cells migrated from OPL to IPL and RNFL + GCL post-ONC.

**Fig. 5.**
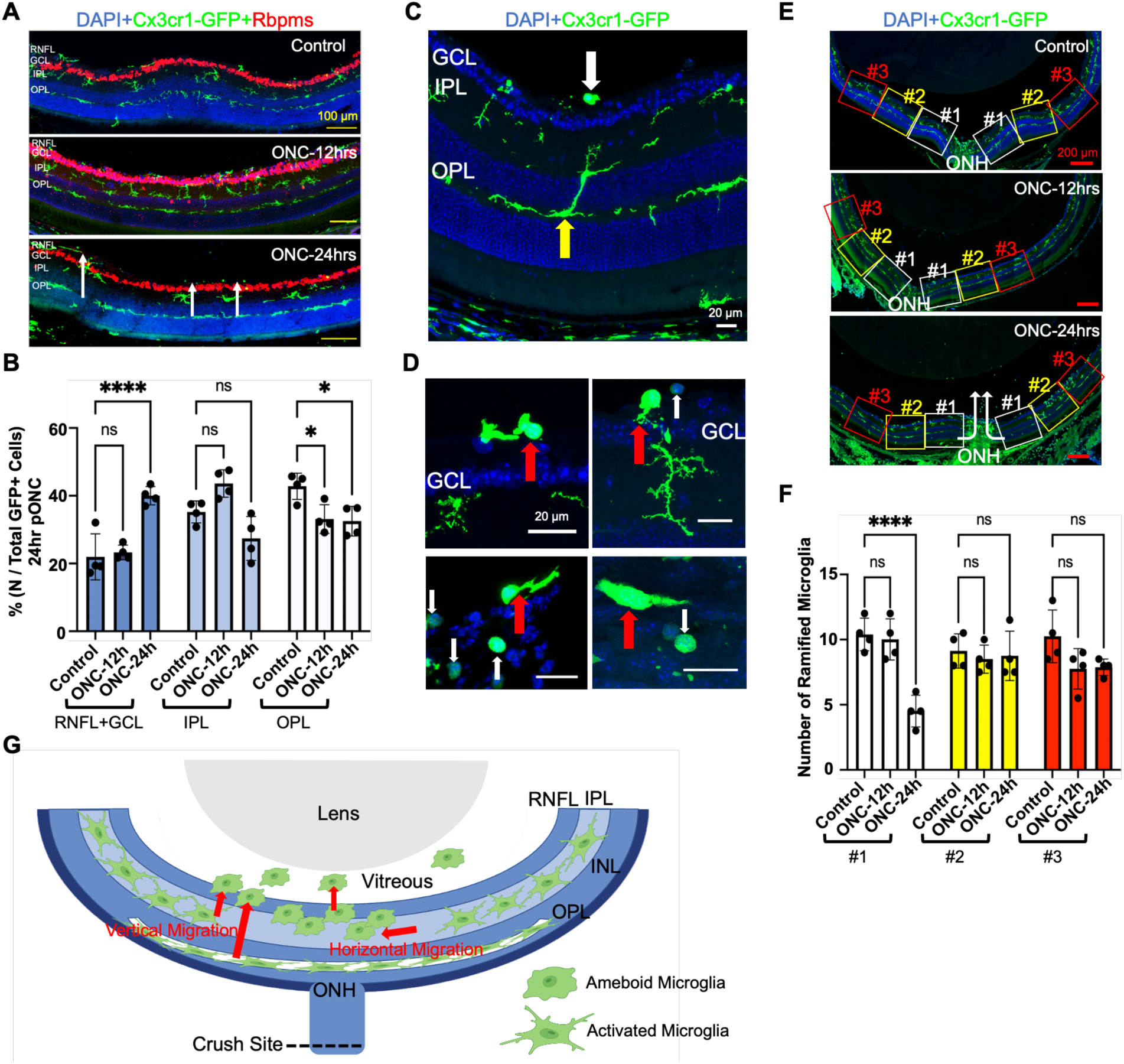
Retinal microglia were mobilized from the outer retina and the vicinity of the ONH. (A) Layer distribution of CX3CR-1+ cells in control and ONC retinas. Rbpms: red. GCL: Ganglion Cell Layer; RNFL: Retinal Nerve Fiber Layer; IPL: Inner Plexiform Layer; OPL: Outer Plexiform Layer. Scale bar: 100 μm. (B) The retinal layer distribution of GFP+ cells in control, ONC-12 h, and ONC-24 h retinas. n = 4 mice. *: p < 0.05; ****: p < 0.0001 (Two-way ANOVA/Bonferroni’s post-hoc test). (C) Microglia captured when migrating from OPL to IPL (yellow arrow). The white arrow indicates amoeboid microglia. (D) Microglia transited from the ramified to amoeboid shape. Red arrows indicate transition state microglia; white arrows indicate amoeboid microglia. (E) Area quantification of ramified microglia in the retina. Each rectangle is 0.1 mm^2^. White arrows indicate the migration direction of retinal microglia. (F) Number of ramified microglia in each area (#1, #2, #3). n = 4 mice. ****: p < 0.0001 (Two-way ANOVA/Bonferroni’s post-hoc test). (G) Illustration of the horizontal and vertical migration of retinal microglia. Red arrows indicate the migration direction of microglia.

Consistent with the above observations, we identified microglia in migration at 12 hours post-ONC. One cell was captured migrating from OPL to IPL, with processes spanning both layers (Fig. 5C). We also observed that some activated microglia in the RNFL + GCL might be in transition to their amoeboid counterparts at 12 hours post-ONC. Some of them are almost amoeboid but have a remnant of processes. Some leave their processes in the IPL while squeezing the cell bodies into the RNFL (Fig. 5D).

Next, we examined the microglial cells in different areas. We found a significant decrease in the number of ramified microglia in the vicinity of the ONH in the ONC-24 h group (4.5 ± 1.2 cells vs 10.4 ± 1.3, n = 4 retinas, p < 1e-4, Fig. 5E, F). Areas further away from the ONH showed no significant changes (Region 2: 8.8 ± 1.9 vs 9.1 ± 1.3 cells, n = 4 retinas, p > 0.99; Region 3: 7.9 ± 0.6 vs 10.3 ± 2.0 cells, n = 4 retinas, p = 0.26). Together, our findings suggest that retinal microglial cells were mobilized from the outer to the inner retina and depleted from the vicinity of ONH, and these cells crossed the ILM and gave rise to the vitreous signals (Fig. 5G).

Interestingly, we noticed that some amoeboid microglia also existed in the anterior segment (Fig. 6A) and the ciliary body (Fig. 6B) of the ONC eyes. At 12 hours post-ONC, amoeboid microglia were detected near the ciliary body and iris, and Iba1+ cells started to appear in the cornea. At 24 hours post-ONC, more amoeboid microglia showed up in the anterior chamber (Fig. 6A, n = 3 replicates). In other words, the release of amoeboid microglial cells from the retina into the vitreous may represent a mechanism by which the posterior eye communicates with the anterior eye.

**Fig. 6.**
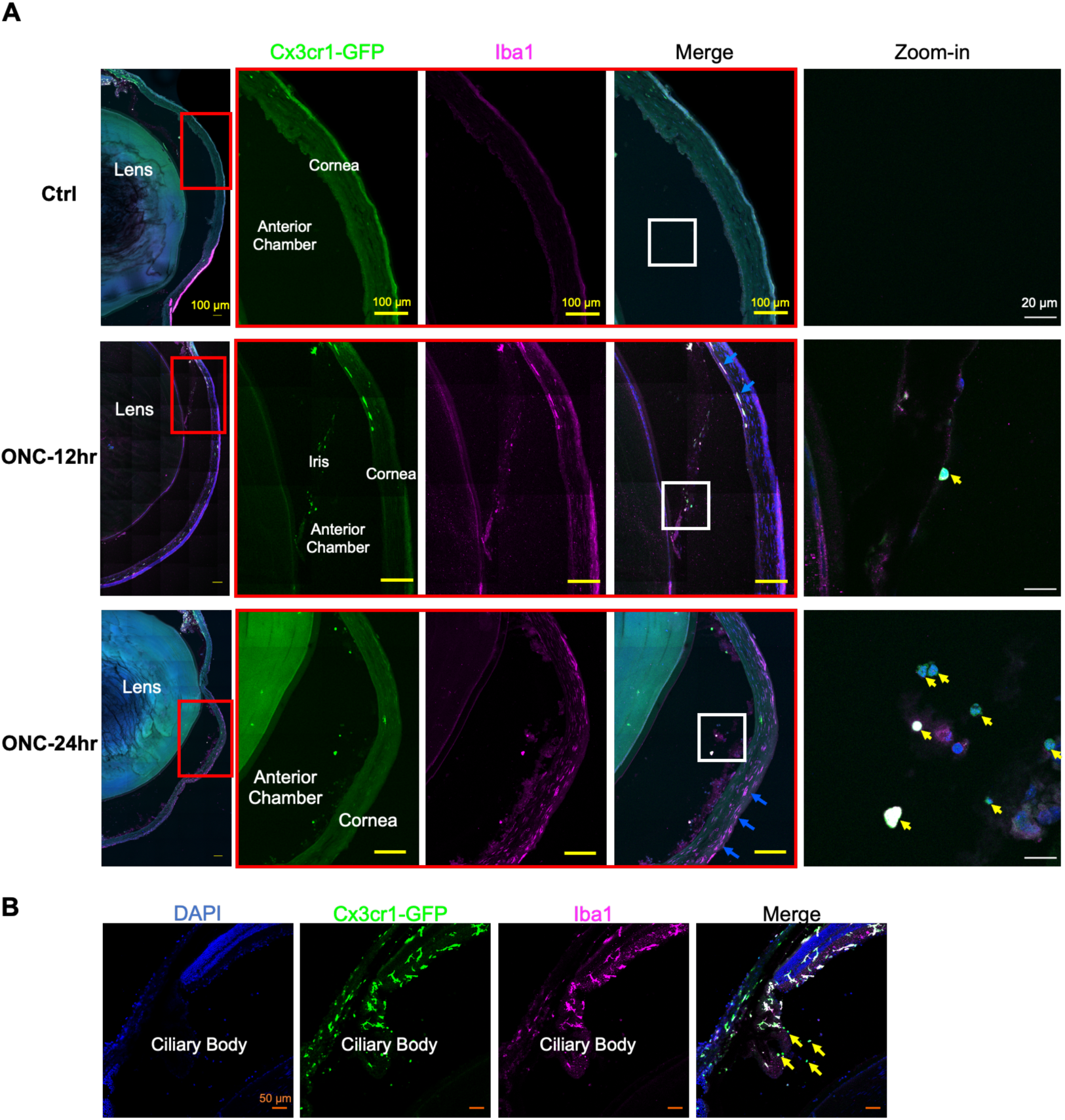
Amoeboid microglia existed in the anterior chamber and ciliary body at 24 hours post-ONC. (A) Microglial cells were immunostained by Iba1 in control, 12 hours, and 24 hours post-ONC CX3CR-1^GFP^ knock-in mice. The red rectangle regions were magnified in the following panels on the right, and further zoomed into the white rectangle. Yellow arrows indicate amoeboid microglia in the anterior chamber; blue arrows indicate Iba1+ cells in the cornea. (B) Amoeboid microglia were detected in the ciliary body and its vicinity (yellow arrows) post-ONC.

### *Interleukin-1*β *(IL-1*β) mRNA was Highly Expressed in Vitreous Amoeboid Microglia

Given the identity of the vitreous cells as activated amoeboid microglia, we next examined the expression of pro-inflammatory cytokines within those cells to specify their molecular properties. We performed RNAscope *in situ* hybridization using *IL-1*β probes in control and 18 hours post-ONC retinas (Fig. 7A). At 18 hours post-ONC, we detected significant *IL-1*β expression in vitreous amoeboid microglia (Fig. 7B). There is a significantly higher percentage of *IL-1*β+ cells among the amoeboid microglia (45.3 ± 12.3 %, n = 4 eyes) compared to the ramified microglia (6.8 ± 6.4 %, n = 4 eyes, p = 1.5e-3, Student’s *t*-test, same below, Fig. 7C). The total number of *IL-1*β+ microglia in the retina also significantly increased at 18 hours post-ONC (9.8 ± 2.6 cells, n = 4 retinas) compared with control (2.8 ± 2.9 cells, n = 4 retinas, p = 1.1e-2, Fig. 7D). Note that an *IL-1*β+ microglia is defined as CX3CR-1+ cells co-labeled with 5 or more *IL-1*β puncta.

**Fig. 7.**
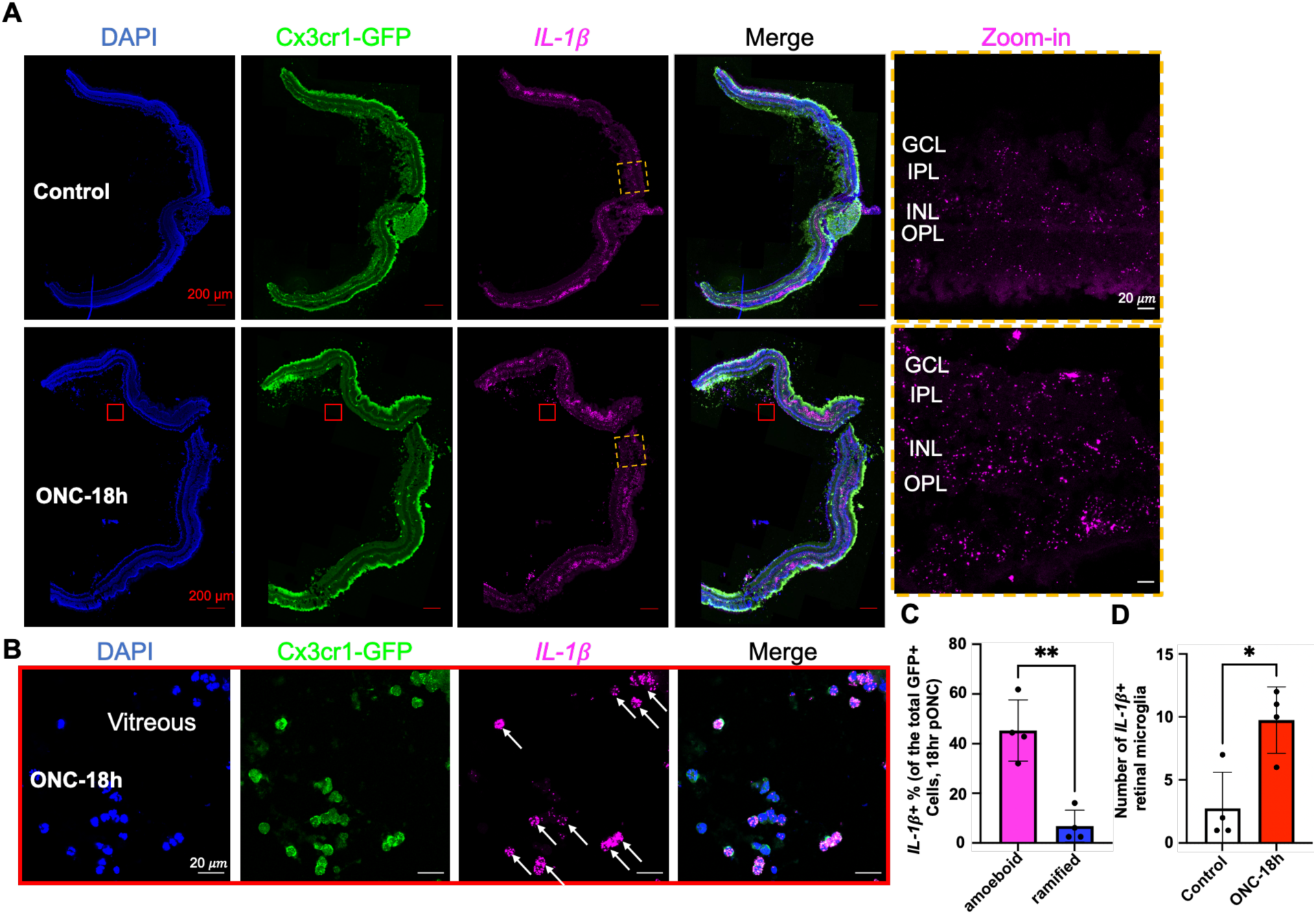
*IL-1β* mRNA was expressed in vitreous amoeboid microglia and elevated in the retina at 18 hours post-ONC. (A) RNAscope staining of *IL-1β* in control and 18 hours post-ONC CX3CR-1^GFP^ knock-in mouse eyes. The orange dashed rectangles were magnified in the zoom-in panel. (B) High magnification images of the panel A vitreous region in red squares. White arrows indicate *IL-1β*-expressing cells. (C) Quantification of the percentage of *IL-1β*+ cells among total amoeboid microglia and total ramified microglia. n = 4 mice. **: p < 0.01 (Student’s *t*-test). (D) The total number of *IL-1β*+ retinal microglial cells in control and 18 hours post-ONC retinas. n = 4 mice. *: p < 0.05 (Student’s *t*-test).

In addition, we examined one of the downstream signaling molecules of the IL-1 pathway: *TNFα*, and detected its increased expression in vitreous amoeboid microglia at 18 hours post-ONC (S-Fig. 3A). At 3 days post-ONC, both expression of *IL-1*β and *TNFα* continued to be elevated in the retina (S-Fig. 3B).

### Interleukin-1 Receptor Antagonist (IL-1Ra) Inhibited the Vitreous Signals and Improved RGC Survival

Interleukin-1 receptor type 1 (IL-1R1) is the primary signaling receptor for IL-1α and IL-1β molecules [52]. To test whether targeting the IL-1R1 can modulate the VHRFs and how it affects RGC survival, we applied an IL-1R1 antagonist called Anakinra right after the ONC surgery. Anakinra is a recombinant form of human IL-1Ra and an FDA-approved drug for rheumatoid arthritis [53]. Here, we tested whether Anakinra inhibited retinal and vitreal inflammation which in turn promotes RGC survival following ONC injury.

First, we tested the feasibility of different delivery methods of Anakinra. We found intravitreal injections induced VHRFs, possibly due to direct damage to the vitreous, while intracameral injection and topical administration did not (S-Fig. 4A). As baseline control, we showed that Anakinra topical treatment did not have significant effects on RGC survival (3487 ± 171 vs 3619 ± 170 cells/mm^2^, n = 6 retinas, p = 0.21, Student’s *t*-test, S-Fig. 4B, C).

Next, we performed ONC on both eyes of the same mouse, with one eye receiving an intracameral injection of Anakinra, and the other eye receiving an equal volume of PBS immediately after surgery. We then assessed the VHRF density at 18 hours post-ONC. Significant decreases in VHRF density were detected in ONC plus Anakinra injection groups (ONC + Ana 40 μg: 309.7 ± 113.8 VHRFs/mm^3^, n = 9 eyes, p < 1e-4; ONC + Ana 20 μg: 667.0 ± 410.4 VHRFs/mm^3^, n = 5 eyes, p = 1.7e-3, One-way ANOVA/Dunnett’s post-hoc test), compared to the ONC plus PBS group (1428 ± 528.5 VHRFs/mm^3^, n = 6 eyes, Fig. 8A, B).

**Fig. 8.**
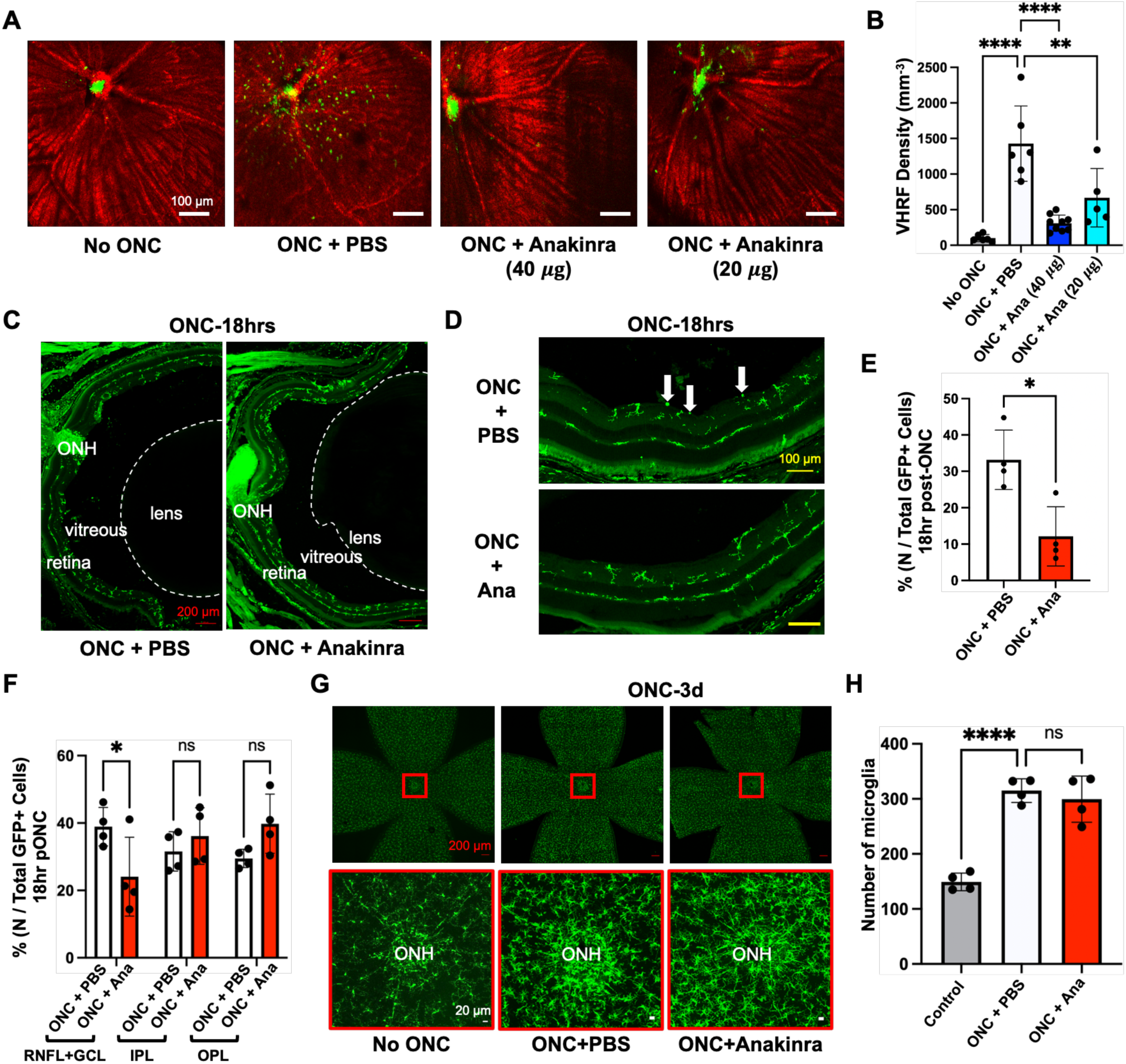
IL-1Ra Anakinra decreased the amoeboid microglia number and inhibited microglia migration, leading to reduced vitreous signals. (A) Vis-OCT images of VHRFs with different drug treatments. (B) Quantification of VHRF density with different drug treatments. **: p < 0.01, ****: p < 0.0001 (One-way ANOVA/Dunnett’s post-hoc test). (C-D) Whole-eye sections of CX3CR-1^GFP^ knock-in mice at 18 hours post-ONC with PBS or Anakinra treatment. White arrows indicate amoeboid microglia. (E) Percentage of amoeboid microglia among all GFP+ microglial cells at 18 hours post-ONC with PBS or Anakinra treatment. n = 4 mice. *: p < 0.05 (Student’s *t*-test). (F) Percentage of microglia within 3 retinal layers with and without Anakinra treatment. n = 4 mice. *: p < 0.05 (Two-way ANOVA/Bonferroni’s post-hoc test). (G) Retina flat mounts of CX3CR-1^GFP^ knock-in mice, no ONC or 3 days post-ONC with PBS or Anakinra treatment. Red squares indicate the zoom-in areas around the ONH (639 μm × 639 μm area). (H) Quantification of microglial cell number within the zoom-in areas of panel F. n = 4 mice. ****: p < 0.0001 (One-way ANOVA/Dunnett’s post-hoc test).

To validate the *in vivo* imaging results, we performed whole-eye sectioning with Anakinra (40 μg) versus PBS post-ONC in CX3CR-1^GFP^ knock-in mice (Fig. 8C, D). We found that the percentage of amoeboid microglia significantly decreased in the ONC plus Anakinra group (12.1 ± 8.1 %, n = 4 retinas) compared to control eyes (33.2 ± 8.2 %, n = 4 retinas, p = 1.2e-2, Student’s *t*-test, Fig. 8E). The percentage of microglia in the RNFL+GCL layer also significantly decreased (24.1 ± 11.7 % vs 38.9 ± 5.7 %, n = 4 retinas, p = 4.2e-2, Two-way ANOVA/Bonferroni’s post-hoc test, Fig. 8F), suggesting inhibited microglia migration to the inner retina. However, the total number of microglia in a 639 μm * 639 μm area around the ONH remained the same between the PBS and Anakinra groups (315.0 ± 21.7 vs 299.5 ± 42.0 cells, n = 4 retinas, p = 0.54, One-way ANOVA/Dunnett’s post-hoc test, Fig. 8G, H). Together, these findings suggested that IL-1Ra (Anakinra) inhibited vitreous signals primarily by reducing the number of amoeboid microglia and inhibiting their migration, without affecting the total microglial population in the retina.

We next compared the RGC survival with Anakinra versus PBS injections at 3 days and 7 days post-ONC. At 3 days post-ONC, significant increases in RGC density were detected in both 40 μg (4009 ± 216 cells/mm^2^, n = 4 retinas, p = 1.8e-3) and 20 μg Anakinra injection groups (3987 ± 236 cells/mm^2^, n = 6 retinas) compared to the ONC plus PBS group (3505 ± 164 cells/mm^2^, n = 7 retinas, p = 9e-4, One-way ANOVA/Dunnett’s post-hoc test, same below, Fig. 9A, B). The ONC plus PBS group survival is consistent with our previous 3 days post-ONC data (3475 ± 343 cells/mm^2^, n = 9 retinas, percentage of difference = 0.86%) [19]. At 7 days post-ONC, there was no significant increase in 40 μg (2684 ± 410 cells/mm^2^, n = 6 retinas, p = 6.2e-2) and 20 μg Anakinra injection groups (2390 ± 217 cells/mm^2^, n = 4 retinas, p = 0.77) compared to the ONC plus PBS group (2230 ± 302 cells/mm^2^, n = 5 retinas, Fig. 9C, D). These results showed that Anakinra delayed RGC loss at 3 days following ONC.

**Fig. 9.**
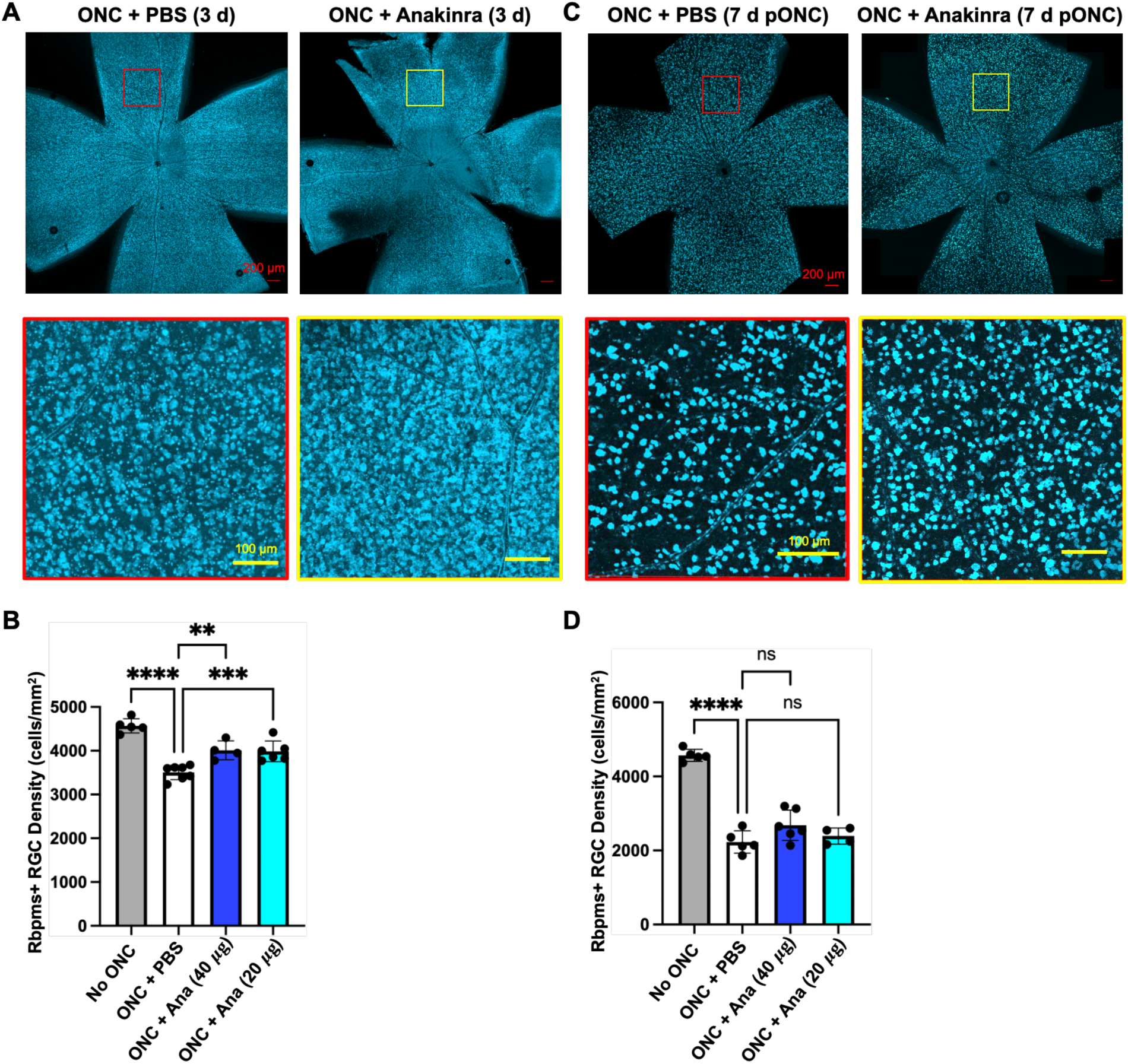
IL-1Ra Anakinra improved RGC survival. (A) Confocal images of flat-mounted retinas immunostained by Rbpms with and without Anakinra treatment at 3 days post-ONC. The bottom panel is the zoom-in image of the top panel. (B) Rbpms+ RGC density with PBS and different dosages of Anakinra treatment at 3 days post-ONC. **: p < 0.01, ***: p < 0.001, ****: p < 0.0001 (One-way ANOVA/Dunnett’s post-hoc test). (C) Confocal images of flat-mounted retinas immunostained by Rbpms with and without Anakinra treatment at 7 days post-ONC. The bottom panel is the zoom-in image of the top panel. (D) Rbpms+ RGC density with PBS and different dosages of Anakinra treatment at 7 days post-ONC. ****: p < 0.0001 (One-way ANOVA/Dunnett’s post-hoc test).

## Discussion

In this study, we investigated the timeline, identity, motility, and functional impact of vitreous hyperreflective foci (VHRFs) in the mouse ONC model. First, we found the VHRFs peaked at 12-24 hours post-ONC, preceding significant RGC loss. Second, we identified the VHRFs as activated amoeboid microglial cells and characterized their migration from the deeper retina to the superficial layer, from the vicinity of the ONH to the ONH/ vitreous. These cells were found in the anterior chamber as well. Finally, *IL-1*β expression was found to be prominent in the vitreous amoeboid microglial cells, and pharmacological inhibition of the VHRFs through IL-1R antagonism significantly improved RGC survival at 3 days post-injury.

### Early indicators of RGC loss

The structural, functional, and molecular signs of RGC degeneration preceding cell death have been investigated. For example, retinal nerve fiber layer (RNFL) and ganglion cell complex (GCC) thinning were detected by near-infrared OCT (NIR-OCT) imaging as indicators of RGC health [54,55]. However, both mouse and human studies have shown that RNFL and GCC thickness measurements lack sufficient sensitivity to detect early neuronal loss [19,56,57]. For functional assessment, pattern electroretinogram (PERG) could detect reductions in RGC response amplitude, serving as an indicator of RGC dysfunction and impending loss [58]. Significant amplitude decrease can be detected at 3 days post-ONC [59]. However, PERG is a less direct measurement, and other diseases, including diabetic retinopathy, Parkinson’s disease, and macular degeneration, also depress PERG, which can cause a false positive diagnosis [60]. Mitochondrial dysfunction and early axonal transport deficits are early events happening within hours of post-disease insults [61]. However, these changes are so subtle that detection often requires invasive *in vivo* imaging or ultrastructural *ex vivo* imaging [62,63].

In our study, the VHRFs detected by *in vivo* vis-OCT imaging after ONC might serve as the earliest warning signal for the entire eye. Our vis-OCT imaging data showed slight increases in VHRFs at 6 hours, and significant increases at 12 hours post-ONC. At 12 hours, our confocal data also showed a few amoeboid microglia reaching the anterior eye, and more cells joined at 24 hours (Fig. 6). These findings suggested that the VHRFs migrated to the vitreous and might serve as an early danger signal for the whole eye. Interestingly, in a rat hypertensive model, VHRFs were identified and histologically identified as “hyalocyte-like Iba1+ cells” [64]. These cells closely resemble our vitreous amoeboid microglial cells in the ONC model in morphology, size, and immunoreactivity, and are very likely the same cells.

In the ONC model, the time course of RGC loss is rapid. In our study, the first significant RGC loss happened at 1 day, with 10 seconds of crush time (Fig. 2). A previous study showed first significant RGC loss occurred at 3 days in both ONC and optic nerve transection (ONT) models using Brn3a RGC marker [65]. Another study showed similar results with the Rbpms marker, while the crush time is 5 seconds instead of 10 [66]. Despite variations depending on crush duration, applied force, and different RGC markers, the same experimenter using the same condition can still be self-consistent. In glaucoma models, in contrast, RGC loss is a slow and progressive process. Common glaucoma models include experimental models, such as laser trabecular photocoagulation [67–69], microbead injection [70], and genetic models, such as DBA/2J [71] and Angiopoietin-1 conditional knockout (A1-cKO) mice [50,72,73], all involving the elevation of IOP due to outflow blockage of the aqueous humor. In the laser photocoagulation model, significant RGC loss was first observed at 1 month post-surgery, with 9.1% cell death [69]. In the microbead injection model, 38.9% of RGC loss was detected at 1 month post-surgery [74]. In DBA/2J mice, significant RGC loss was not detected until 10 months [75], and in Angiopoietin-1 conditional knockout mice, significant RGC loss was detected at 3 months [50]. Importantly, at 24 hours, VHRFs peaked before the earliest significant RGC death was detected (Fig. 2). We also detected a low density of VHRFs under chronic IOP elevation in A1-cKO mice. Although VHRFs were present, their density was markedly lower than in ONC, consistent with previous reports [64]. Long-term tracking of the VHRFs and RGC loss with chronic IOP elevation might be needed in the future.

### Neuroinflammation and Neuroprotection

Neuroinflammation has been extensively studied in different eye diseases. For example, one of the main causes of diabetic retinopathy (DR) is the activation of glial cells, including astrocytes, Müller cells, and microglia [76]. In DR, microglia undergo proliferation, migration, and changes in their morphology (from ramified to amoeboid), accompanied by up-regulation of pro-inflammatory factors including IL-1β, TNFα, IL-6, IL-8, and IL-18, contributing to chronic retinal inflammation, blood–retina barrier (BRB) breakdown, and vascular leakage [77]. In age-related macular degeneration (AMD), macrophages, dendritic cells (DCs), and neutrophils stimulate T cells and B cells and participate in choroid neovascularization (CNV). Cytokines, including IL-1β, TNFα, IL-6, IL-8, IL-10, IL-17, TGF-β, IFN-γ, etc, could promote angiogenesis and thus diminish retinal pigment epithelium (RPE) cells, photoreceptors, and choroidal capillaries [78]. In glaucoma, retinal microglia, astrocytes, and Müller glia were activated under elevated intraocular pressure (IOP) [79,80], with up-regulation of TNFα, IL-1α, IL-1β, IFNγ, IL-6, and MIP-1α, etc. [81]

Retinal glial cells affect RGC survival by phagocytosis, glutamate clearance, neurotropic factor maintenance, and inflammatory cytokine production [82]. Microglia, among others, also play a complex role in RGC survival post-damage. On the one hand, they produce pro-inflammatory cytokines and inducible nitric oxide synthase (iNOS), which induce RGC apoptosis and tissue damage [40]. On the other hand, their phagocytic effects and release of neurotrophic factors also promote RGC survival [83,84]. Therefore, it is always essential to accurately target the detrimental aspects of microglia while keeping their beneficial traits. For example, whole-body depletion of microglia by dietary CSF1R antagonist has no effect or even detrimental effect on RGC survival post-ONC or glaucomatous insult [85–87]. It was suggested that microglia whole-body depletion upregulated the neurotoxic A1 genes in astrocytes, thus jeopardizing RGC survival [86]. In contrast, selectively targeting pro-inflammatory pathways or molecules, including NLRP3 inflammasome knockout [31], IL-1 receptor antagonism [46], IL-6 knockout [47], and IL-1α/ TNFα/ C1q triple knockout [48], substantially preserves RGC survival. These results align with the findings of our current study that specifically inhibiting the pro-inflammatory IL-1 receptor pathway is protective to RGCs post-damage. In our study, IL-1 receptor antagonism inhibited the morphology change and migration of microglia cells at early timepoints post-ONC. However, it remains to be investigated whether IL-1 receptor antagonism directly or indirectly affects these processes, since loss of amoeboid morphology is likely to impede microglial migration as well.

Anakinra, the interleukin-1 receptor antagonist (IL-1Ra) we applied in this study, is an FDA-approved medication used to treat a series of immune-mediated disorders, including rheumatoid arthritis, cryopyrin-associated periodic syndromes, familial Mediterranean fever, and Still’s disease [88]. Anakinra is a recombinant human IL-1Ra that blocks the biological activity of IL-1 by competitively inhibiting its binding to the IL-1 type I receptor (IL-1RI) [89]. It has a molecular weight of 17257.6 Da, which likely explains the ineffectiveness of topical treatment (S-Fig. 4B, C), as molecules larger than 500 Da are difficult to cross the cornea [90]. In ocular diseases, Anakinra has been applied in clinical trials for dry eye disease [91] and uveitis [92]. In a uveitis mouse model, subcutaneous injection of Anakinra significantly reduced inflammatory cells in the vitreous and retina [93]. Moreover, in a blast-mediated traumatic brain injury model, Anakinra treatment reduced Müller glia activation and increased survival of axon bundles in the optic nerve [46]. These findings are consistent with our reduced VHRFs and improved RGC survival in the ONC model following Anakinra injection.

Overall, this work identified a novel early *in vivo* danger sign for RGC loss and highlighted the potential of targeting pro-inflammatory signaling at early stages of injury as a feasible neuroprotective strategy. Future studies will be needed to determine whether VHRFs serve as early indicators of RGC loss in chronic glaucoma patients, and whether IL-1 receptor antagonism provides neuroprotection in this context. Addressing these questions will be critical for translating this novel biomarker, along with associated inflammation-targeting therapies, into clinical applications.

## Acknowledgments

We thank Dr. Marta Grannonico and Dr. Mingna Liu for their technical support. In particular, we thank Yujuan Yi for her assistance with RNAscope. We also thank Prof. Ignacio Provencio for his insightful discussions. This study was supported by the National Institutes of Health Grant R01EY035088 (X.L. and H.Y.), the Knights Templar Eye Foundation Career-starter Research Grant (J.W.), and the BrightFocus Foundation National Glaucoma Research Award (X.L.).

## Notes

### Competing Interest Statement

The authors have declared no competing interest.

