## Supplemental Figures for "Vitreous Inflammation as an Early Indicator of Retinal Ganglion Cell Loss Following Acute Optic Nerve Injury in Mice"

#### Supplemental Figure Legends

**S-Fig. 1 The VHRFs originated from the optic nerve head (ONH) and dispersed over time.** (A) Quantification illustration of the VHRF pattern within the same eye at different time points. By drawing concentric circles of 200, 300, and 400 μm radius from the ONH, the vitreous *en face* images were divided into 3 areas, and the number of dots was quantified in each area. (B) Change of inflammatory signal number with radius at 12, 24, and 72 hours post-ONC. n = 5 mice. *: $p < 0.05$; ***: $p < 0.001$ (Two-way ANOVA/Bonferroni's post-hoc test).

**S-Fig. 2 More ameboid microglia were the M1 pro-inflammatory subtype; a proportion of the amoeboid microglia were undergoing proliferation.** (A) CD11c and CD206 staining for M1 and M2 microglia subtypes CX3CR-$1^{GFP}$ knock-in mouse retinas. Violet and red arrows indicate CD11c+ and CD206+ cells, respectively. (B) Percentage of CD11c+ and CD206+ cells among total CX3CR-$1^{GFP}$+ amoeboid microglia. n = 4 mice (Student's *t*-test). (C) A vitreous amoeboid microglial cell captured during cell division (red arrow). (D) Ki67 staining for proliferating cells in CX3CR-$1^{GFP}$ knock-in mouse retina.

**S-Fig. 3 RNAscope of *TNFα* in control and ONC retinas.** (A) *TNFα* is expressed in amoeboid microglia at 18 hours post-ONC. Yellow arrows indicate *TNFα*-expressing cells. (B) *TNFα* and *IL-1β* expression at ONC-18 hours vs ONC-3 days retinas. Red squares are magnified in the zoom-in panel.

**S-Fig. 4 Feasibility test of different delivery methods for IL-1Ra Anakinra.** (A) Vis-OCT images of the VHRFs post different treatment conditions: no treatment, 24 hours post-ONC, 24 hours post-PBS intravitreal injection, 24 hours post-PBS anterior chamber injection, and Anakinra topical administration. (B) Confocal images of flat-mounted retinas immunostained by Rbpms with and without daily topical Anakinra treatment (5 μL/eye, 20 μg/μL) at 3 days post-ONC. (C) Rbpms+ RGC density with PBS and Anakinra treatment at 3 days post-ONC. n = 6 mice. (Student's *t*-test).

### S-Fig. 1

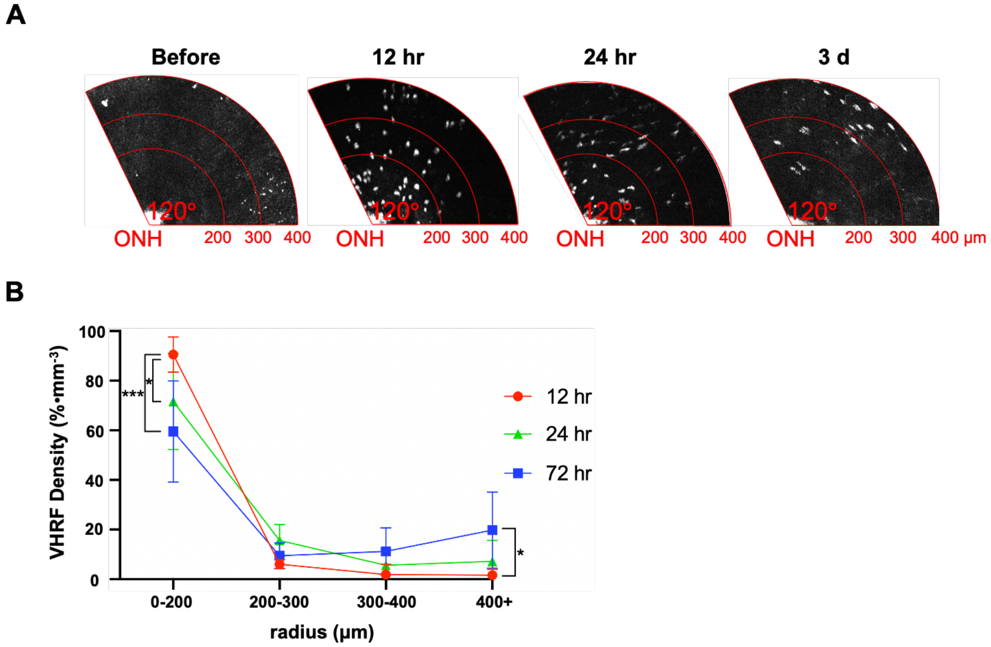

A
Before
12 hr
24 hr
3 d
120°
ONH
200 300 400
400 µm
B
VHRF Density (%•mm⁻³)
100
80
60
40
20
0
***
*
0-200
200-300
300-400
400+
radius (µm)
12 hr
24 hr
72 hr

### S-Fig. 2

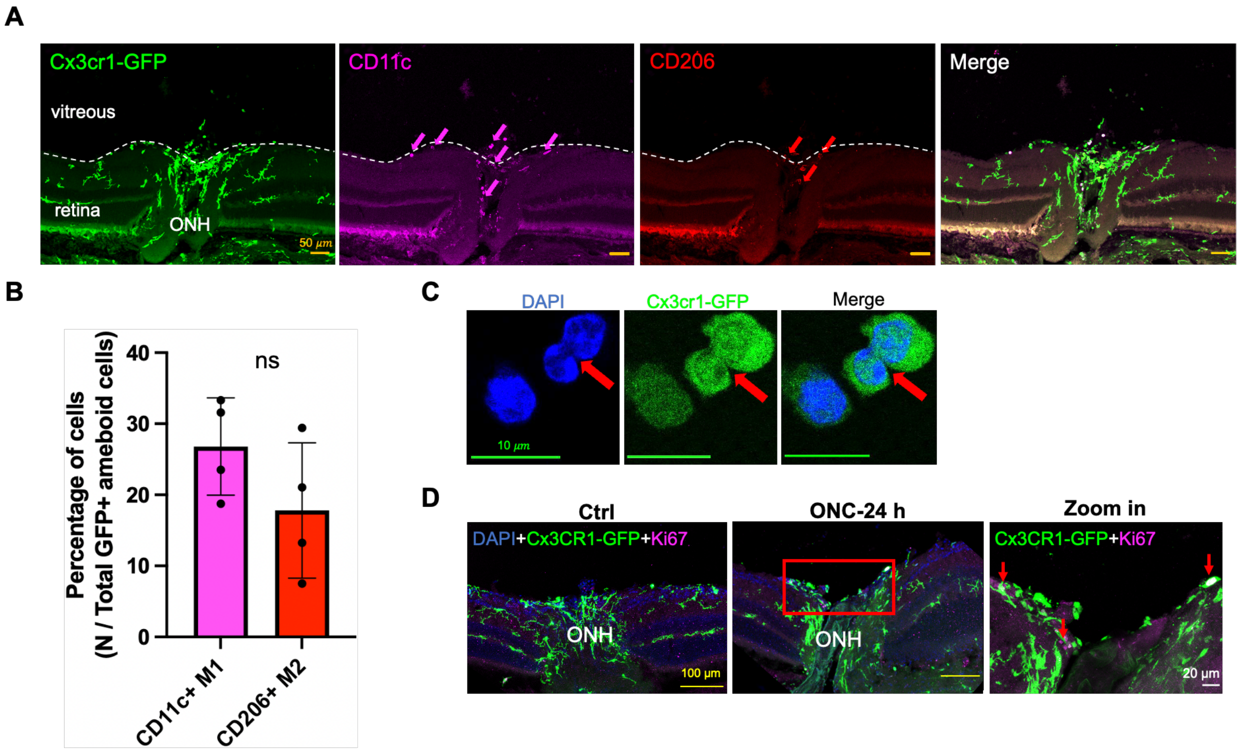
A
Cx3cr1-GFP
vitreous
retina
ONH
50 μm
CD11c
CD206
Merge
B
Percentage of cells
(N / Total GFP+ ameboid cells)
40
30
20
10
0
ns
CD11c+ M1
CD206+ M2
C
DAPI
Cx3cr1-GFP
Merge
10 μm
D
Ctrl
ONC-24 h
Zoom in
DAPI+Cx3CR1-GFP+Ki67
ONH
100 μm
Cx3CR1-GFP+Ki67
20 μm

### S-Fig. 3

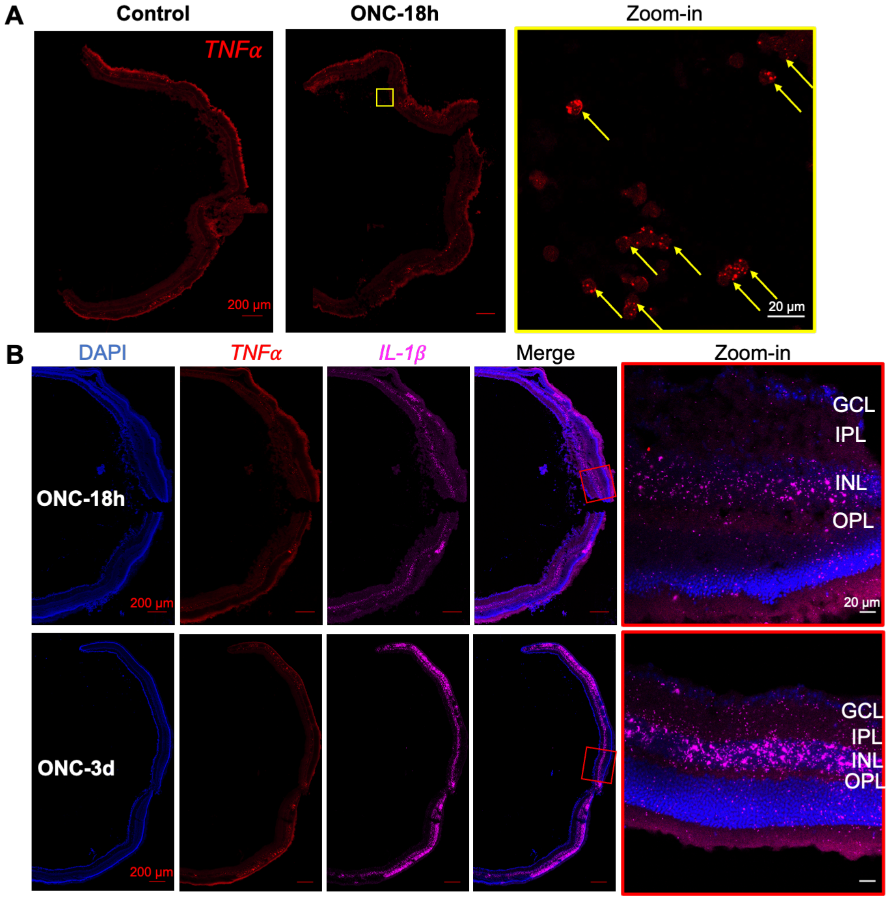
A
Control
ONC-18h
Zoom-in
TNFα
200 μm
20 μm
B
DAPI
TNFα
IL-1β
Merge
Zoom-in
ONC-18h
200 μm
GCL
IPL
INL
OPL
20 μm
ONC-3d
200 μm
GCL
IPL
INL
OPL

### S-Fig. 4

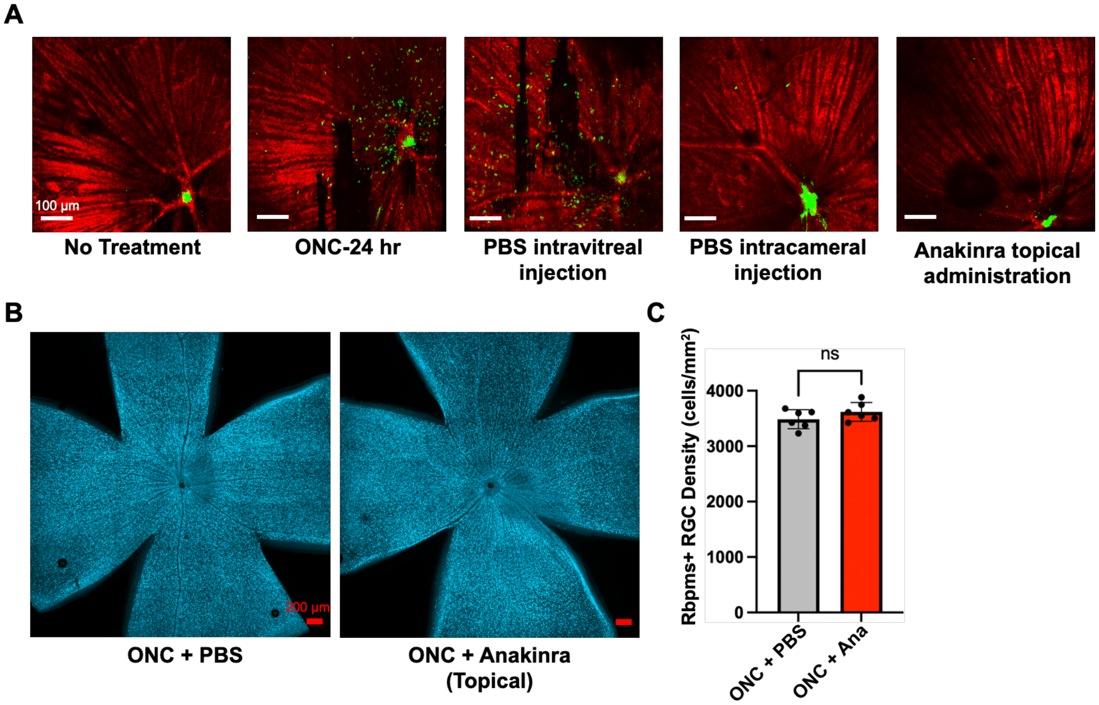
A
100 µm
No Treatment
ONC-24 hr
PBS intravitreal injection
PBS intracameral injection
Anakinra topical administration
B
200 µm
ONC + PBS
ONC + Anakinra (Topical)
C
Rbpms+ RGC Density (cells/mm²)
ns
4000
3000
2000
1000
0
ONC + PBS
ONC + Ana
